# HIF1α controls steroidogenesis under acute hypoxic stress

**DOI:** 10.1101/2024.08.05.606625

**Authors:** Stephen Ariyeloye, Deepika Watts, Mangesh Jaykar, Cagdas Ermis, Anja Krüger, Denise Kaden, Barbara K. Stepien, Vasileia Ismini Alexaki, Mirko Peitzsch, Nicole Bechmann, Ali El-Armouche, Ben Wielockx

## Abstract

**Background:** Hypoxia is a critical physiological and pathological condition known to influence various cellular processes, including steroidogenesis. While previous studies, including our own, have highlighted the regulatory effects of Hypoxia-Inducible Factor 1α (HIF1α) on steroid production, the specific molecular mechanisms remain poorly understood. This study investigates the role of hypoxia and HIF1α in steroid biosynthesis across multiple experimental models during acute exposure to low oxygen levels.

**Methods:** To assess the extent to which acute hypoxia modulates steroidogenesis, we employed several approaches, including the Y1 adrenocortical cell line, an *ex vivo* adrenal gland explant model, and a conditional HIF1α-deficient mouse line in the adrenal cortex. We focused on various regulatory patterns that may critically suppress steroidogenesis.

**Results:** In Y1 cells and adrenal gland explants, hypoxia induced the upregulation of specific microRNAs, leading to the suppression of mRNA levels of key steroidogenic enzymes and reduced steroid hormone production. The hypoxia/HIF1α-dependent induction of these microRNAs and the consequent modulation of steroid production were confirmed in vivo. Notably, using our conditional HIF1α-deficient mouse line, we found that the increase in miRNA expression under hypoxic conditions is directly dependent on HIF1α. Furthermore, the regulation of steroidogenic enzymes (e.g., StAR and Cyp11a1) and steroid production occurred at the level of protein translation, revealing an unexpected layer of control under hypoxic conditions *in vivo*.

**Conclusions:** These findings elucidate the molecular mechanisms underlying acute hypoxia-induced changes in steroid biosynthesis and may also be useful in developing new strategies for various steroid hormone pathologies.

## Background

The hypoxia signaling pathway is essential for cellular and systemic adaptations to reduced oxygen levels and for the modulation of many pathophysiological processes (1). Regulation of this critical system is primarily orchestrated by the activities of the hypoxia-inducible factor (HIFα) subunits, which are in turn tightly regulated by the HIF prolyl hydroxylase domain proteins (PHDs), the von Hippel-Lindau tumor suppressor protein (VHL), and the factor inhibiting HIF1 (FIH) under physiological oxygen concentrations, leading to HIFα inactivation and rapid degradation (2, 3). However, under reduced oxygen levels, the functionality of these regulators is hampered, which activates HIFα signaling (4). One of the pivotal processes regulated by the hypoxia/HIF pathway is steroidogenesis in which steroid hormones are synthesized by the catalytic conversion of cholesterol (5). Steroid hormones play a critical role in controlling several important developmental and physiological processes and any disruption in their production can lead to various health problems, including reproductive disorders, obesity, and hypertension (6).

We, along with others, have previously described a significant role for HIFs in the production of steroids (7–11); specifically, in a recent study, we revealed a critical function for HIF1α in regulating the synthesis of virtually all steroids in the mouse adrenal gland under steady-state conditions. Additionally, a lack of HIF1α in the adrenocortical cells resulted in a sustained increase in the expression of steroidogenic enzymes, which in turn boosted steroid production. Conversely, HIF1α stabilization in these cells markedly decreased steroidogenesis, as evidenced by a significant reduction in the transcription of the enzymes involved in steroid catalysis and a consequent decrease in steroid output. While available literature also indicates that the hypoxia/HIF pathway can hinder steroid production, the precise molecular mechanisms by which this pathway affects steroidogenesis in acute situations are not well understood (12).

MicroRNAs (miRNAs/miRs) are a class of non-coding ribonucleic acids (ncRNAs) that are recognized for their role in the negative regulation of gene expression (13). These molecules function primarily by binding to complementary sequences within the 3’ untranslated regions (3’UTR) of their target messenger RNAs (mRNAs), resulting in the repression of gene expression either by interfering with translation or by facilitating the degradation of the target mRNA (14). MicroRNAs are pivotal in controlling a wide array of biological processes, such as angiogenesis, apoptosis, hematopoiesis, and growth and differentiation of cells (12, 15, 16). Furthermore, alterations in miRNA activity have been linked to various diseases including cancer, diabetes, metabolic and cardiovascular disorders (17). Under hypoxic conditions, the expression of a specific subset of microRNAs, known as hypoxia-induced microRNAs or hypoxaMIRs, is modulated (18, 19). These hypoxia-regulated microRNAs are critical for modulating cellular responses to low oxygen levels and ensuring proper adaptation (20). Additionally, miRNAs are known to play a regulatory role in steroid production, with several studies highlighting their impact on the regulation of genes involved in steroidogenesis (21, 22). Thus, while available literature indicates that a complex interplay between hypoxia/HIFs and miRNAs influences steroidogenesis (23, 24), a complete understanding of this crosstalk in the regulation of steroidogenesis is still lacking.

Hence, we aimed to elucidate the molecular pathways by which hypoxia adversely affects steroidogenesis by exposing multiple steroidogenic models to acute hypoxic conditions. Results from both a cell line model and an *ex vivo* adrenal gland model demonstrate that hypoxia critically suppresses steroidogenesis, which is associated with the direct upregulation of a class of miRNAs targeting a set of key enzymes involved in steroid production. *In vivo* mouse models showed that this increase in miRNA expression under hypoxic conditions is directly dependent on HIF1α. Confirming studies in conditional HIF1α-deficient mice revealed that modulation of this process during the acute phase of hypoxic stress is primarily mediated by a HIF1α-dependent regulation of StAR and Cyp11a1 translation, revealing an elaborate layer of regulatory complexity.

## Methods

### Cell culture

Murine Y1 adrenocortical cells were cultured in Dulbeccós Modified Eaglès Medium (DMEM) supplemented with 10% Fetal Calf Serum (FCS), 2.5% horse serum, and 1% penicillin-streptomycin, at 37°C in a humidified incubator with 5% CO_2._ DMEM was additionally supplemented with 2.5% UltroSerG (15950-017, Pall Life Sciences) and 1% Insulin-Transferrin-Selenium during experiments. For hypoxia experiments, cells were exposed to 5% O_2_ levels in a 37°C incubator with 5% CO_2_ (Whitley H35 Hypoxystation) (Don Whitley Scientific Limited, United Kingdom). Dimethyloxaloylglycine (DMOG, Biomol) was used at 1 mM.

### *Ex vivo* culture of adrenal glands

Mouse adrenal glands were cultured *ex vivo* at 21% or 5% oxygen levels in DMEM containing 2.5% Nu serum (355100, Corning), 1% penicillin-streptomycin and 1% Insulin-transferrin-selenium and maintained at 37°C in a humidified incubator with 5% CO_2_. Hypoxia treatment of the adrenal glands was conducted in the hypoxia chamber (Whitley H35 Hypoxystation) (Don Whitley Scientific Limited, United Kingdom).

### Mice

Mice used in this study were bred and housed under specific pathogen-free (SPF) conditions at the Experimental Centre of the Medical Theoretical Center (MTZ, Technical University of Dresden-University Hospital Carl-Gustav Carus, Dresden, Germany. The Akr1b7:cre-*Phd2/Hif1^ff^*/*^ff^*(P2H1^Ad.Cortex^) mouse line was generated by crossing Akr1b7:cre mice to *Phd2^f/f^* and *Hif1α^f/f^* as described elsewhere (7). All mice (both genders - including WT) used in this study were bred on a C57BL/6J background (backcrossed at least 10 times) and pups were born at normal Mendelian ratios. The primers used to genotype transgenic mice have been described previously (7). The isolated adrenal glands were either used for *ex vivo* experiments or snap frozen in liquid nitrogen and stored at − 80 °C for gene expression or hormone analysis. Breeding of all mice and animal experiments were in accordance with local guidelines on animal welfare and were approved by the Landesdirektion Sachsen, Germany.

### Hypoxia treatment of mice

Wild-type mice were challenged with 9% O_2_ for 24 hours, and adrenal glands were collected, snap-frozen, and stored at −80 °C. To achieve hypoxia (9% O_2_), mice were placed in a BioSpherix chamber. Oxygen pressure within the chamber was consistently sensed using a ProOx P110 compact oxygen controller (Parish, New York) and adjusted according to the set point by nitrogen infusion. Mice were placed in the chamber in standard cages containing bedding and supplied with food and water *ad libitum*.

### Steroid hormone measurement

Steroids hormones in cell culture supernatants, mouse plasma and adrenal glands were measured as previously described (25, 26).

### Western Blotting

For Western blot, adrenal tissue samples were prepared as previously described (26) and protein concentrations were measured using the bicinchoninic acid assay (BCA) (Thermo Fisher). Briefly, 20 µg of protein were separated on a 4-12% Bis-Tris protein gels (Thermo Fisher) and transferred. The membranes were blocked (5% milk) and incubated with primary antibodies against StAR (∼28 kDa, 1:1000 – Cat. #8449) and Cyp11a1 (∼55 kDa, 1:1000 – Cat. #14217) or Metavinculin (∼145 kDa, 1:1000 – Cat. #18799) (all Cell Signaling) overnight at 4°C. The following day, membranes were washed and incubated with secondary rabbit IgG HRP (R&D Systems) for 1 hour at room temperature. The membranes were then exposed to Fusion Fx (peqlab, VWR). Quantification was performed using Fiji (ImageJ distribution 1.52K) (27).

### mRNA real-time PCR

RNA was isolated from adrenal glands or Y1 cells using the RNA Easy Plus micro kit (Qiagen) (Cat. # 74034, Qiagen), the RNA Easy Plus mini kit (Cat. # 74134, Qiagen) or the PARIS™ Kit (Thermo Fisher Scientific). Reverse transcription was carried out with the iScript cDNA Synthesis Kit (BIO-RAD, Feldkirchen, Germany). Gene expression analysis was performed by quantitative real-time PCR with the ‘Ssofast Evagreen Supermix’ (BIO-RAD, Feldkirchen, Germany). Real-Time PCR Detection System-CFX384 (BIO-RAD, Feldkirchen, Germany) was used for quantification of synthesized cDNA. All mRNA expression levels were calculated relatively to beta-2 microglobulin (*B2m*) or eukaryotic translation elongation factor 2 (*Eef2*) housekeeping genes using the 2(-ddCt) method, where ddCT was calculated by subtracting the average control dCT (e.g., 21% O_2_ for Y1 cells, adrenal gland *ex vivo*, WT or P2H1^Ad.Cortex^), from dCT of all samples individually. Primer sequences used are described in supplementary material (Supplementary Table 1).

### MicroRNA real-time PCR

RNA was isolated from adrenal glands or Y1 cells using the RNA Easy Plus micro kit (Cat. # 74034, Qiagen), the RNA Easy Plus mini kit (Cat. # 74134, Qiagen) or the PARIS™ Kit (Thermo Fisher Scientific). Reverse transcription was carried out using the Mir-X™ miRNA First Strand Synthesis Kit (Takara Bio). The TB Green kit (Takara Bio) was used for detecting miRNA expression levels. Relative gene expression was determined using the 2(−ddCt) method with U6 as the housekeeping gene. Primer sequences used are described in supplementary material (Supplementary Table 1).

### Bioinformatic analysis

The miRDB (www.mirdb.org) and TargetScan (www.targetscan.org) computational algorithms were used to identify predicted targets of microRNAs. MicroRNA sequences were determined using the miRPathDB 2.0 (www.mpd.bioinf.uni-sb.de), Genescript (www.genscript.com) and MirGeneDB (www.mirgenedb.org) databases, as well as from published reports (28–35).

### Statistical analyses

All data are presented as mean ± SEM. For assessing the statistical significance of two experimental groups, a two-tailed Mann–Whitney U-test or unpaired t-test with Welch’s correction (after testing for normality with the F test) was used, unless stated otherwise in the text. For assessing the statistical significance of multiple experimental groups with one variable, the 1-way ANOVA t-test followed by the Tukey’s post hoc test was used. Statistical differences presented in the figures were considered significant at p-values below 0.05. All statistical analyses were performed using the GraphPad Prism v10.02 for Windows or higher (GraphPad Software, La Jolla California USA, www.graphpad.com).

## Results

### Hypoxia induces miRNA expression and represses steroidogenesis in Y1 cells

Several studies have documented the impact of hypoxia on steroid production across various steroidogenic models (7–9). Yet, the precise molecular pathways through which hypoxia and HIF1 suppress steroidogenic genes remain incompletely understood, although prior research suggests that HIF1 may primarily play an indirect role (36, 37). Given the significant role of miRNAs as epigenetic regulators that can negatively influence gene expression, and the proposed interaction between HIFs and miRNAs in mediating gene suppression (5), we aimed to explore the effects of hypoxia on miRNA expression; specifically those targeting steroidogenic enzymes. First, through bioinformatic analysis and literature search (detailed in the Materials & Methods section), we identified a number of microRNAs predicted to target steroidogenic enzymes. We chose to study 5 miRNAs that target the *Star* mRNA because of its primary role in catalyzing the rate-limiting step of steroid production (38). Previous studies have reported the influence of some of these miRNAs on the expression of steroid biosynthetic enzymes (21, 39). An overview of these identified microRNAs presumed to suppress steroidogenic enzymes, including their predicted targets, are displayed in Figure 1A.

**Figure 1.**
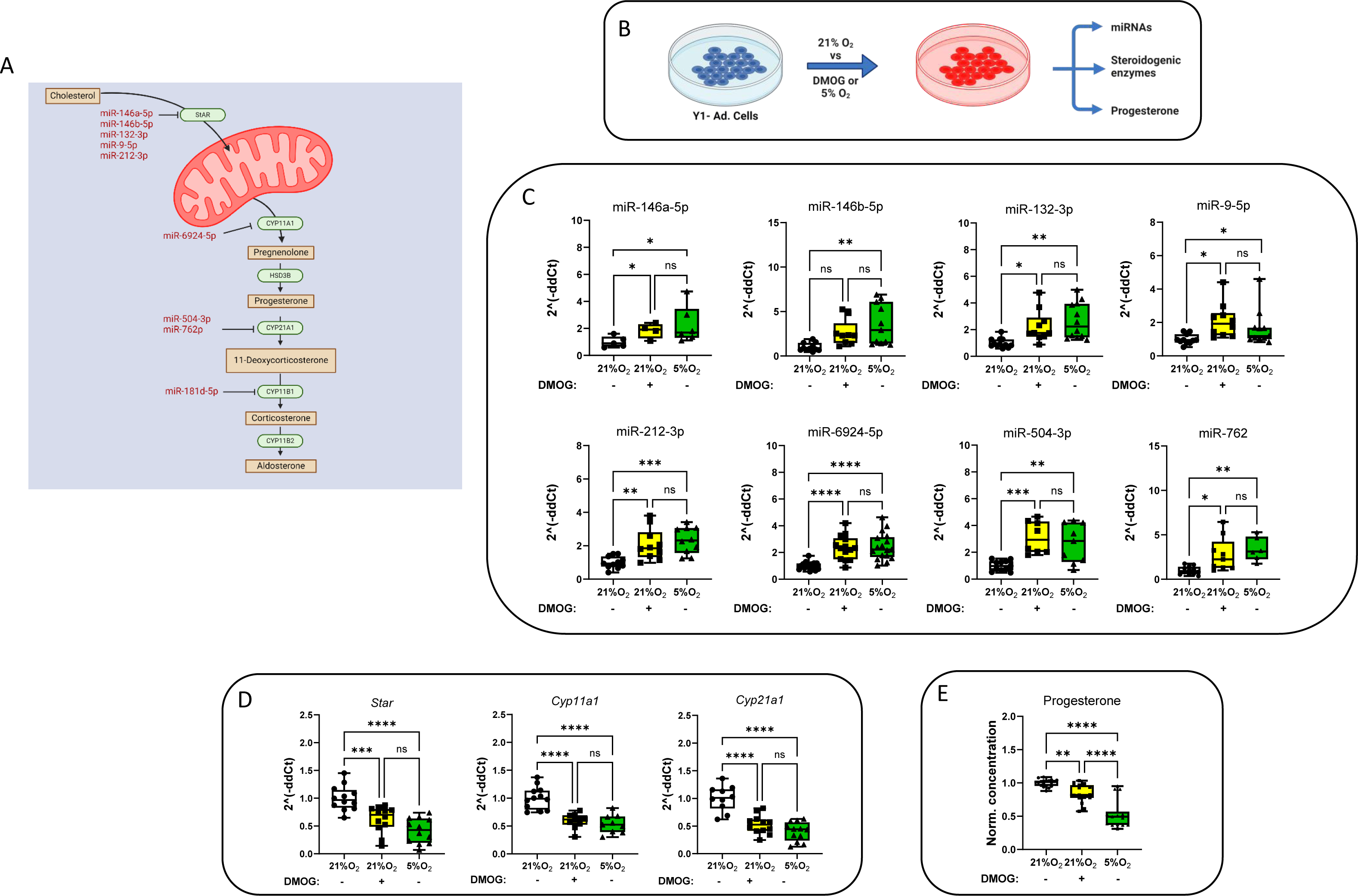
Hypoxia induces microRNA expression and represses steroidogenesis in Y1 cells. **A.** Schematic representation of the identified microRNAs putatively repressing steroidogenic enzymes; miR-146a-5p, miR-146b-5p, miR-132-3p, miR-9-5p, and miR-212-3p target *star*, miR-6924-5p targets *cyp11A1*, miR-504-3p and miR-762 target *cyp21a1*, miR-181d-5p targets c*yp11b1*. **B.** Schematic representation of the experimental set-up. **C.** Real-time qPCR analysis showing a general upregulation of the expression of the identified microRNAs in Y1 cells cultured under hypoxic conditions (5% O_2_ or DMOG) compared to the normoxic group (n= 4-17). **D.** qPCR-based mRNA expression analysis of key steroidogenic enzymes in Y1 cells cultured under hypoxic conditions (5% O_2_ or DMOG) compared to the normoxic group (n= 9-12). **E.** Reduction in progesterone secretion in Y1 cells exposed to hypoxic conditions (5% O_2_ or DMOG) compared to the normoxic group (n= 15-18). Values are expressed as mean ± the SEM (*p < 0.05, **p < 0.01, ***p < 0.001, ****p < 0.0001). One-way ANOVA with the post hoc Tukey’s test. P values of miR-146a-5p and miR-9-5p were calculated using a one-tailed Mann– Whitney U-test. All data were normalized to the normoxic group (21% O_2_) average value.

Next, to ascertain responsiveness to hypoxia, we evaluated the transcriptional activity of these miRNAs in Y1 cells under hypoxic conditions by exposing them to low oxygen (5% O_2_) or treating them with the hypoxia mimetic DMOG for 24h compared to the cells from normoxic (21% O_2_) conditions. (Figure 1B). DMOG stabilizes HIFs even in the presence of high oxygen levels due to blocking PHD enzyme activity thereby mimicking the molecular effects of hypoxia. Compared to normoxic control samples, selected miRNAs showed an overall upregulation in the expression, both under hypoxia and with DMOG (Figure 1C). To further investigate the impact of these hypoxia-induced microRNAs on their target steroidogenic enzymes, we measured mRNA levels of the key enzymes in Y1 cells and found a dramatic reduction in all mRNA levels measured (Figure 1D). Notably, this reduction was associated with a decrease in progesterone, the only measurable steroid in this setting (Figure 1E). Taken together, these observations support the notion that hypoxia plays a critical role in suppressing steroidogenesis by enhancing miRNA expression.

### Hypoxia-enhanced microRNA expression associates with reduced steroidogenesis in adrenal glands cultured *ex vivo*

Acknowledging the limitations inherent to the Y1 cell model, particularly their deficient expression of specific steroidogenic enzymes and steroids (40), we sought to validate our findings within a more physiologically relevant context. Hence, we used adrenal glands isolated from WT mice and cultured them *ex vivo* under low oxygen (5% O_2_) or atmospheric conditions for 28 hours. This approach allowed us to closely examine the expression profiles of miRNA and steroid catalyzing enzymes, apart from quantifying steroid production levels in a controlled hypoxic environment. (Figure 2A).

**Figure 2.**
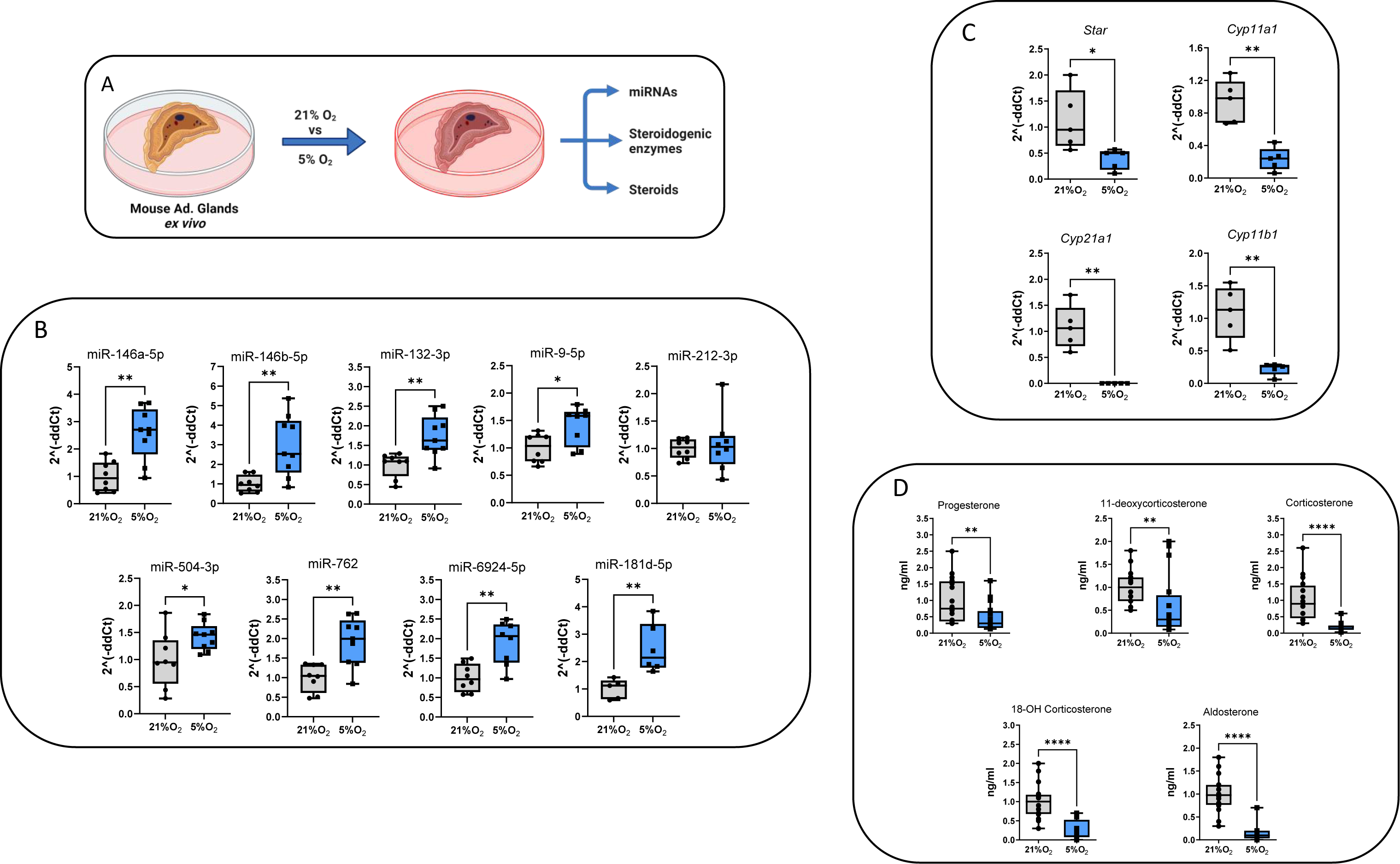
Hypoxia induction of microRNA expression correlates with reduced steroidogenesis in *ex vivo* cultured adrenal glands. **A.** Schematic representation of the *ex vivo* experimental set-up. **B.** Real-time qPCR results demonstrating upregulation of the entire collection of identified microRNAs (except for miR-212-3p) in adrenal glands cultured *ex vivo* under low oxygen pressure (5% O_2_) (n = 8-9). **C** Real-time qPCR analysis showing a reduction in the expression of *star, cyp21a1, cyp11a1* and *cyp11b1* steroidogenic enzymes (n = 5). **D.** A significant decrease in the steroid levels in *ex vivo* cultured adrenal glands under low oxygen pressure (5% O_2_), including corticosterone and aldosterone (n = 16). Values are represented as the mean ± SEM (*p < 0.05, **p < 0.01, ***p < 0.001, ****p < 0.0001) Mann–Whitney U-test. All data were normalized to the normoxic group (21% O_2_) average value.

Hypoxic conditions upregulate the expression of all tested miRNAs (except for miR-212-3p) (Figure 2B). This upregulation at least parallels the expression patterns observed in Y1 cells under hypoxic conditions, and suggests consistent hypoxia-responsive behavior of these miRNAs across different experimental conditions. Concurrently, quantitative PCR analysis revealed a reduction in the expression of key steroidogenic enzymes (Figure 2C), coupled with a significant decrease in steroid production levels, including that of corticosterone and aldosterone (Figure 2D). These findings not only confirm our initial observations made in Y1 cells, but in a broader context suggest that hypoxia-induced miRNAs influence steroidogenesis at the cell and also adrenal gland level.

### Diverse responses to hypoxia-induced miRNA expression and steroidogenesis *in vivo*

We next aimed to explore the dynamics of miRNA expression and activity, and its effects on steroidogenesis under a more complex systemic condition. Therefore, we expanded our investigation to an *in vivo* setting by acutely exposing WT mice to a reduced oxygen environment (9% O_2_) for 24 hours. Readouts from these mice were compared with those from littermate controls maintained at atmospheric conditions (Figure 3A). Analysis of blood cell content in circulation revealed that this acute exposure to hypoxia was too short to substantially increase red blood cell (RBC) formation although it reduced myeloid cell numbers and enhanced platelets (Supplementary Figure 1A). In line with our cell culture and *ex vivo* findings, *in vivo* studies revealed a uniform increase in the levels of miRNAs (except for miR-181d-5p) and confirming their sensitivity to acute hypoxic stress (Figure 3B). Intriguingly, despite broad induction of miRNAs, their impact on the mRNA expression of their corresponding steroidogenic enzymes presented a complex pattern as mRNA levels of 3 out of 4 key enzymes were significantly increased, contrasting with the general expectation of miRNA-mediated suppression (Figure 3C). Therefore, we performed western blot analyses for StAR and Cyp11a1 enzymes and found that, despite the significant increases in mRNA levels, their protein levels remained unchanged (Figure 3D and Supplementary Figure 1B), suggesting hypoxia-dependent interference during the translation process. In accordance with these results, steroid levels in the adrenal glands remained largely unchanged, and notably, aldosterone levels were significantly diminished, highlighting the intricate regulatory effects induced by acute hypoxia in a living mouse (Figure 3E).

**Figure 3.**
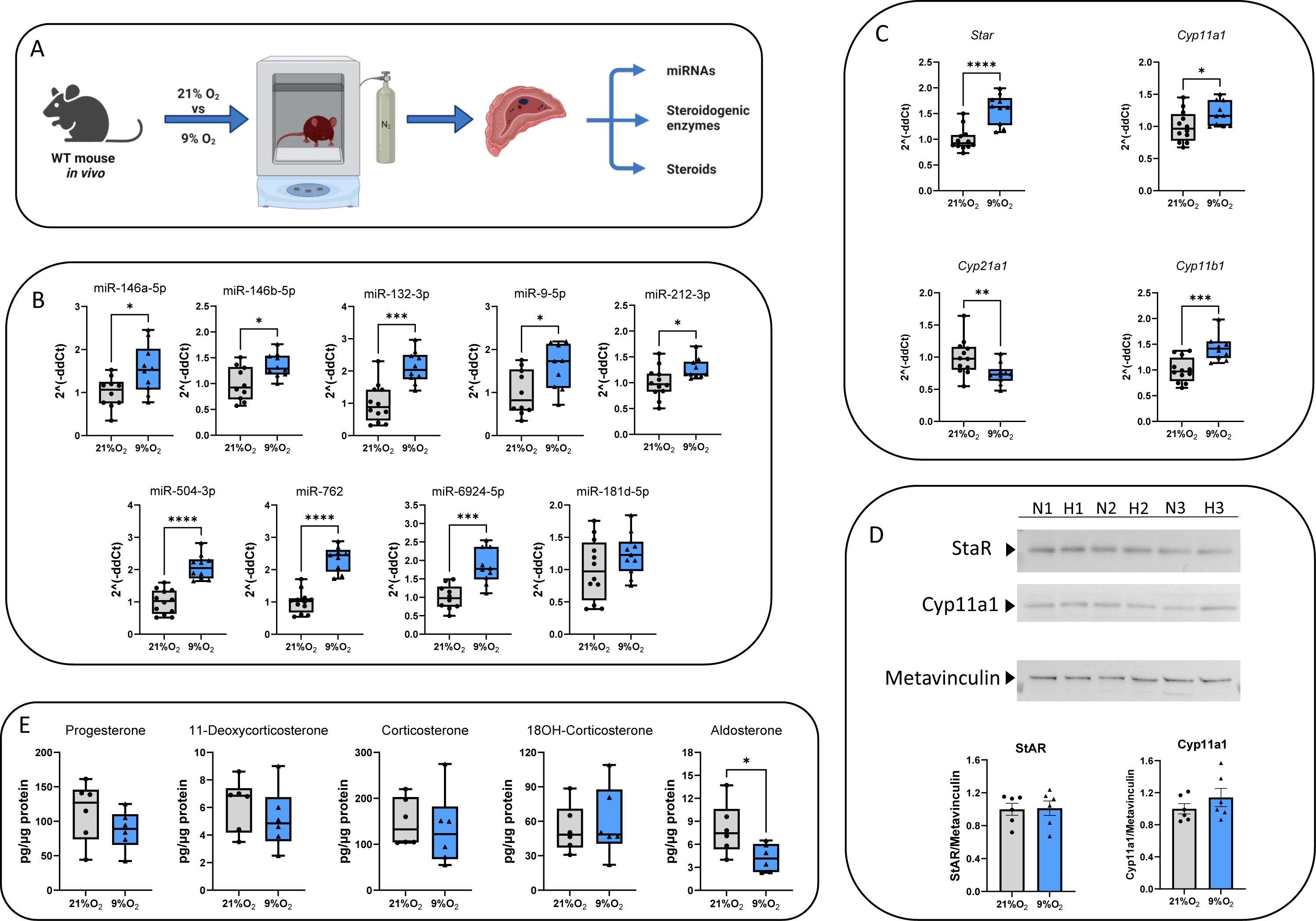
Acute hypoxia regulated steroidogenesis in WT mice *in vivo*. **A.** Schematic representation of the experimental set-up. **B.** Real-time qPCR results demonstrating a uniform increase in the expression levels of the identified microRNAs (except for miR-181d-5p) in the adrenal glands of mice exposed to low oxygen pressures (9% O_2_) for 24 hours compared to the normoxic group (n =10-12). **C.** Real-Time qPCR analysis of the steroidogenic enzyme expression in the adrenal glands of mice exposed to low oxygen pressures (9% O_2_) for 24 hours compared to the normoxic group (n=10-12). **D.** StAR and Cyp11A1 protein levels were analyzed via Western blot using Metavinculin as a loading control. All protein levels were quantified using FIJI and normalized to Metavinculin levels. N = 21% O_2_, H = 9% O_2_ (representative blot with 6 individual samples). **E.** Steroid levels in the adrenal glands of mice exposed to hypoxic stress (9% O_2_) for 24 hours did not experience a corresponding increase in the first 24 hours of exposure compared to the normoxic group (n =6). Values are represented as the mean ± SEM (*p < 0.05, **p < 0.01, ***p < 0.001, ****p < 0.0001) Mann–Whitney U-test. All data in B, C and D were normalized to the normoxic group (21% O_2_) average value.

### *In vivo* HIF1α mediates miRNA expression independent from steroidogenesis during acute hypoxia

To investigate at what level HIF1α might be involved in acute hypoxia-induced steroidogenesis, we used our previously described mouse line in which HIF1α is deficient in adrenocortical cells, whilst the activity of HIF2α is increased already under normoxic conditions (Akr1b7:cre-*Phd2/Hif1^ff^*/*^ff^*(P2H1^Ad.Cortex^)) (7). These P2H1^Ad.Cortex^ mice were subjected to acute hypoxia (9% O_2_ for 24h) and compared with mice of the same genotype under atmospheric conditions (Figure 4A). Firstly, we showed that the blood cell pattern in these mice was similar to that in the experiment with WT mice (Supplementary Figure 2A). However, hypoxic exposure of these P2H1^Ad.Cortex^ mice displayed no changes in expression levels of 8 out of 9 microRNAs compared to atmospheric conditions, indicating a direct influence of HIF1α in adrenocortical cells on microRNA expression (Figure 4B). Consistently, the mRNA levels of *StAR* and *Cyp11b1* enzymes remained largely unchanged under hypoxic conditions. However, we found a significant decrease in the expression of *Cyp11a1* and *Cyp21a1* (Figure 4C). Contrasting with these mRNA patterns, protein levels of StAR and Cyp11a1 were significantly increased compared to their normoxic counterparts (Figure 4D and Supplementary Figure 2B), aligning with the observed increases in progesterone and corticosterone levels in the adrenal glands after acute hypoxic exposure (Figure 4E).

**Figure 4.**
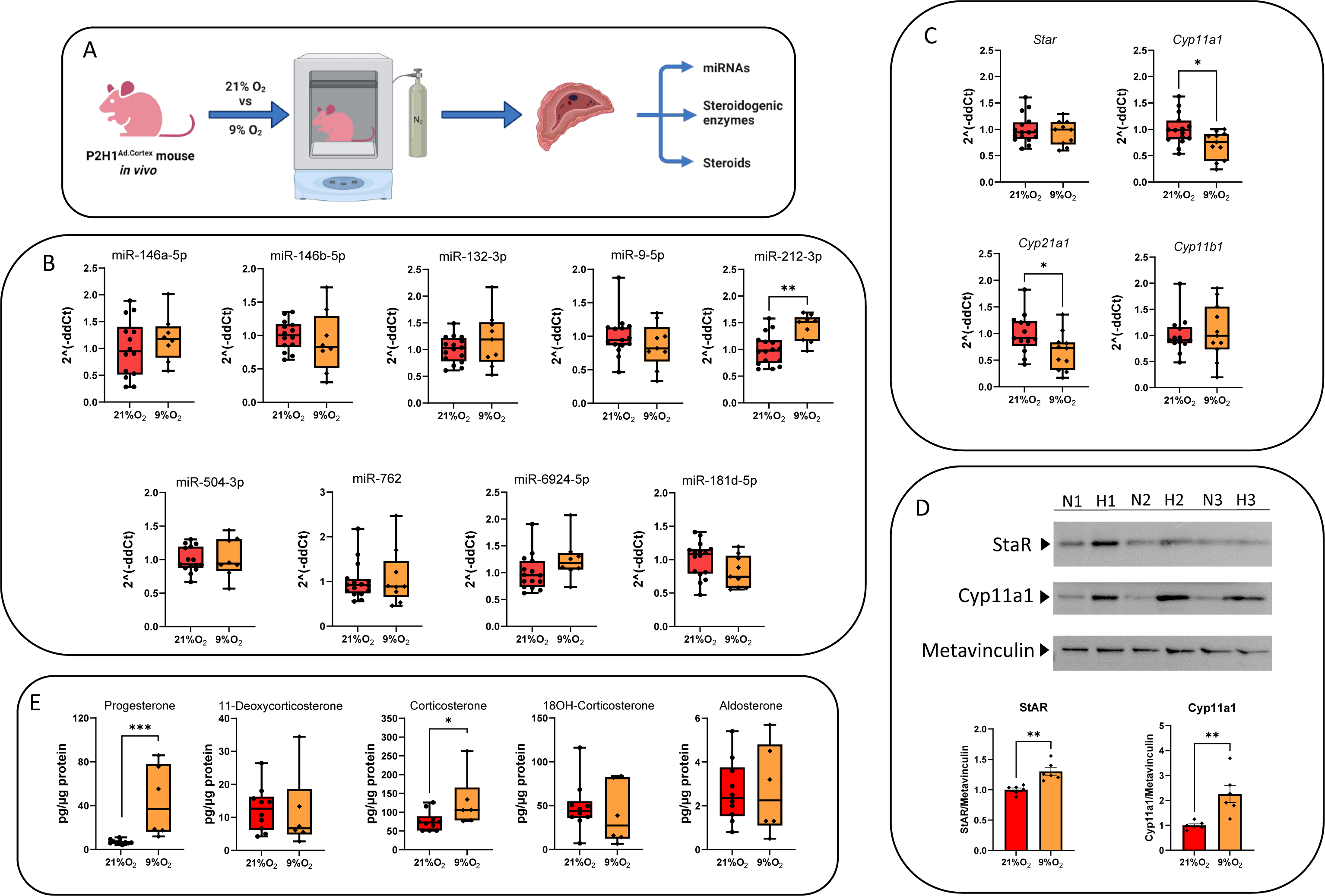
Acute hypoxic modulation of microRNA/steroidogenesis *in vivo* is regulated in a HIF-1α dependent manner. **A.** Schematic representation of the experimental setup. **B.** Real-time PCR analysis shows 8 out of the 9 identified microRNAs displaying no changes in their expression levels under both normoxic and hypoxic conditions in P2H1^Ad.Cortex^ adrenal glands after acute hypoxia (9% O2) (24 hours) compared to their normoxic counterparts (n= 8-16). **C.** Real-time qPCR analysis of the expression levels of *star, cyp21a1,* and *cyp11A1* and *cyp11b1* steroidogenic enzymes in P2H1^Ad.Cortex^ adrenal glands after acute hypoxia (9% O_2_ for 24 hours) compared to their normoxic counterparts (n= 10-16). **D.** StAR and Cyp11A1 protein levels were analyzed via Western blot using Metavinculin as a loading control. All protein levels were quantified using FIJI and normalized to Metavinculin levels. N = 21% O_2_, H = 9% O_2_ (representative blot with 6 individual samples). **E.** Steroid production in the P2H1^Ad.Cortex^ adrenal glands after acute hypoxia (24 hours) compared to their normoxic counterparts (n= 6-10). Values are represented as the mean ± SEM (*p < 0.05, **p < 0.01, ***p < 0.001, ****p < 0.0001) Mann–Whitney U-test. All data in B, C and D were normalized to the normoxic group (21% O_2_) average value.

These *in vivo* experiments prompted further detailed evaluations between the WT and P2H1^Ad.Cortex^ results in order to enhance our understanding of the effects of acute hypoxia and HIF1α on steroidogenesis. When we compared the expression patterns of all microRNAs at 24h after hypoxia, we observed that, except for miR-212-3p, the expression of all other miRNAs was directly associated with the presence of HIF1α (Figure 5A). In addition, we found that the mRNA levels of four key enzymes involved in steroidogenesis were also dependent on HIF1α, although their expression patterns contrasted with those of the microRNAs (Figure 5B). Remarkably, and in contrast to their mRNA levels, both StAR and Cyp11a1 proteins were significantly increased in the cKO mice (Figure 5C), leading to significantly higher end product steroids - progesterone, glucocorticoid and aldosterone (Figure 5D). Taken together, these data suggest that HIF1α is a limiting factor in the ultimate translation of these steroidogenic enzymes, and establish HIF1α as a critical, versatile regulator at multiple stages of steroidogenesis during the first 24 hours of acute hypoxia *in vivo*.

**Figure 5.**
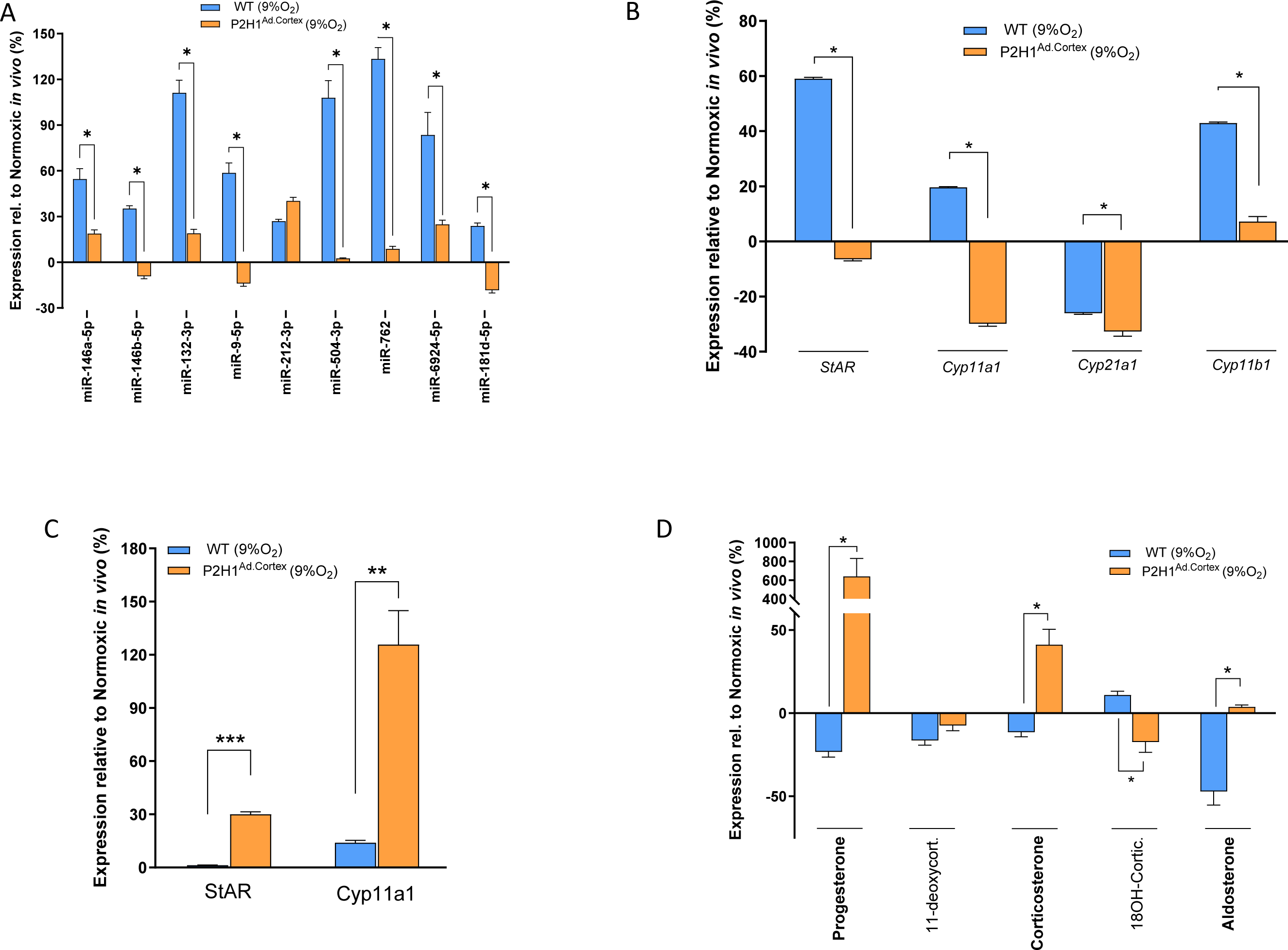
Acute hypoxic and HIF1α regulation of steroidogenesis *in vivo*. **A.** Real-time qPCR analysis of microRNAs from adrenal glands of hypoxia treated WT and P2H1^Ad.Cortex^ mice (24 hours at 9% O_2_) compared to their respective normoxic counterparts (n=10-16). **B.** Gene expression analysis of key steroidogenesis enzymes from adrenal glands of hypoxia treated WT and P2H1^Ad.Cortex^ mice (24 hours at 9% O_2_) compared to their respective normoxic counterparts (n=10-16). **C.** Protein expression analysis of StAR and Cyp11a1 from adrenal glands of hypoxia treated WT and P2H1^Ad.Cortex^ mice (24 hours at 9% O_2_) compared to their respective normoxic counterparts (n=6). **D.** Steroids produced from adrenal glands of hypoxia treated WT and P2H1 mice (24 hours at 9% O2) compared to their respective normoxic counterparts (n=6-10). Values are represented as the mean ± SEM (*p < 0.05, **p < 0.01) Mann–Whitney U-test.

## Discussion

In this study, we explored the acute hypoxic response and its potential repressive effects on steroidogenesis across three different models. Our research demonstrates, for the first time, that the induction of a specific cluster of microRNAs is dependent on hypoxia and HIF1α, which correlates with decreased steroidogenesis in both *in vitro* and *ex vivo* settings. Furthermore, using WT and P2H1^Ad.Cortex^ mice, we identified an additional HIF1α-dependent translational regulation of StAR and Cyp11a1. Collectively, our findings reveal novel insights into how HIF1α regulates steroidogenesis during the acute phase of hypoxia across various cellular, organ, and systemic contexts.

MicroRNAs are known to be potent post-transcriptional regulators of gene expression, and they primarily act by silencing specific target genes by enhancing their degradation (41) or inhibiting their translation (42). Several studies using different model systems have investigated the activities of hypoxia-inducible miRNAs (43, 44), and the role of microRNAs in the modulation of steroidogenesis has been documented (45, 46). Although recent research suggests that hypoxia/HIF may influence steroidogenesis by modulating miRNA expression, comprehensive studies providing a mechanistic understanding of steroidogenesis at the cellular, tissue, and systemic levels have not been performed.

Our study revealed that in response to acute hypoxia, a distinct set of miRNAs targeting at least four steroidogenic enzymes are consistently upregulated across different models. This upregulation is primarily controlled by HIF1α, introducing a novel aspect to our understanding of how hypoxia signaling can intervene very early in the steroidogenesis process. This aligns with existing literature on hypoxia-responsive microRNAs, miR-212 and miR-132, which were previously described to regulate steroidogenesis (21, 39). However, a number of microRNAs used in this study have not yet been implicated in the control of steroidogenesis (47–49). The induction of these miRNAs in our experimental settings is believed to lead to the repression of steroid biosynthesizing enzymes, thereby reducing steroid production —a hypothesis that was confirmed in Y1 cells *in vitro* and *ex vivo* cultured adrenal glands under acute hypoxia. Hypoxia/HIF1α-dependent suppression of gene expression is largely, if not entirely, indirect (36, 37), and our results are consistent with previous studies in steroid-producing human H295R cell lines and the marine medaka ovary (23, 24, 50), emphasizing the indirect impairment of steroid production by hypoxia/HIF1α and miRNAs.

Interestingly, we noted a multifaceted response between hypoxia/HIF1α-induced miRNA expression, enzyme expression patterns and steroidogenesis *in vivo*, highlighting the complex nature of steroidogenesis under acute hypoxic conditions in a complex system. Contrary to the direct correlation between miRNA upregulation and decreased steroid production observed in Y1 cells and *ex vivo* cultured adrenal glands, our findings suggest the involvement of additional HIF1α-dependent regulatory layers *in vivo*. This observation was evident as 24-hour hypoxia resulted in significantly enhanced mRNA levels of the *StAR* and *Cyp11a1* enzymes but not at the protein level. However, in the absence of HIF1α in cortical cells we found increased enzyme expression opposite to their mRNA concentration. At this stage, it is reasonable to speculate that, in response to acute hypoxia in the mouse, steroidogenesis is regulated by an additional HIF1α-associated regulatory system. For instance, it has been described that mammalian target of rapamycin complex 1 (mTORC1), a central mRNA-to-protein translation activator, is inhibited in hypoxia and affects the global translation rate of mRNAs (51). It has also been demostrated that global mRNA-to-protein translation elongation rates can be decreased due to eEF2 inhibition under hypoxic conditions (52). In addition, also miRNAs can bind to the 3’ untranslated regions (UTRs) of their target mRNAs and repress their translation. This interaction can block translation initiation or elongation, leading to reduced protein synthesis while maintaining high mRNA levels (42). More research is therefore warranted to better understand which regulatory mechanisms are at play in the hypoxia-HIF1α-miRNA-steroidogenesis axis, especially in complex biological systems such as a living animal.

Taken together, our data clearly demonstrates that HIF1α negatively regulates steroid production during the acute phase of hypoxia in an adrenocortical cell line, isolated adrenal glands, and mice. How hypoxia/HIF1α specifically affects these different processes during steroidogenesis, and whether the novel set of steroidogenic miRNAs identified here have additional regulatory effects, as has been suggested for mir-132-3p, remains to be unraveled (21). This complex regulatory web highlights the need for further research to elucidate the specific interactions between hypoxia/HIF1α, microRNA dynamics, and steroidogenic pathways that shape the physiological response to low oxygen levels in a nuanced and context-dependent manner.

## Conclusion

Our study demonstrates that HIF1α plays a critical role in modulating steroidogenesis under acute hypoxic conditions, primarily through the induction of specific microRNAs and their regulation of key steroidogenic enzymes. This reveals an unexpected layer of control, offering new insights into the molecular mechanisms of hypoxia-induced changes in steroid biosynthesis.

## Statements & Declarations

### Ethics approval and consent to participate

Breeding of all mice and animal experiments were in accordance with the local guidelines on animal welfare and were approved by the Landesdirektion Sachsen, Germany (TVA12/2019, TVV38/2022, TVvG21/2023).

### Consent for publication

Not applicable

### Availability of data and materials

Data is provided within the manuscript or supplementary information files. Materials are available upon reasonable request

### Authors’ contributions

S.A. designed and performed the majority of experiments, analyzed data, and wrote the manuscript. D.W. designed, performed, and supervised experiments, analyzed data, and contributed to the discussion. M.J., C.E., A.K., D.K., M.P., performed experiments and analyzed data. N.B., B.K.S., V.I.A. and A.E.A. provided tools and contributed to the discussion. B.W. designed, performed, and supervised experiments and the overall study, analyzed data, and wrote the manuscript.

## Acknowledgements

We would like to thank Dr. Vasuprada Iyengar for English Language editing.

---

**Supplementary Figure 1. Sysmex analyses of blood from WT mice**

(A) Number of different blood cells in circulation from WT mice 24 hours after incubation at 21% or 9% O_2_ (#cells/µl). (B) Ponceau staining of the Western blot shown in Figure 3D after transfer and before analyses with the indicated antibodies as well as all uncropped blots of the pictures shown in Figure 3D.

**Supplementary Figure 2. Sysmex analyses of blood from WT mice**

(A) Number of different blood cells in circulation from P2H1^Ad.Cortex^ mice 24 hours after incubation at 21% or 9% O_2_ (#cells/µl). (B) Ponceau staining of the Western blot shown in Figure 4D after transfer and before analyses with the indicated antibodies as well as all uncropped blots of the pictures shown in Figure 4D.

## References

1. Semenza GL. Hypoxia-inducible factor 1 and the molecular physiology of oxygen homeostasis. Journal of Laboratory and Clinical Medicine. 1998;131(3):207–14.

2. Semenza GL. HIF-1 and mechanisms of hypoxia sensing. Curr Opin Cell Biol. 2001;13(2):167–71.

3. Watts D, Gaete D, Rodriguez D, Hoogewijs D, Rauner M, Sormendi S, et al. Hypoxia Pathway Proteins are Master Regulators of Erythropoiesis. Int J Mol Sci. 2020;21(21).

4. Metzen E, Ratcliffe PJ. HIF hydroxylation and cellular oxygen sensing. Biological Chemistry. 2004;385(3-4):223–30.

5. Ariyeloye S, Kammerer S, Klapproth E, Wielockx B, El-Armouche A. Intertwined regulators: hypoxia pathway proteins, microRNAs, and phosphodiesterases in the control of steroidogenesis. Pflugers Arch. 2024.

6. Miller WL, Auchus RJ. The Molecular Biology, Biochemistry, and Physiology of Human Steroidogenesis and Its Disorders. Endocr Rev. 2011;32(1):81–151.

7. Watts D, Stein J, Meneses A, Bechmann N, Neuwirth A, Kaden D, et al. HIF1α is a direct regulator of steroidogenesis in the adrenal gland. Cellular and Molecular Life Sciences. 2021;78(7):3577–90.

8. Marchi D, Santhakumar K, Markham E, Li N, Storbeck KH, Krone N, et al. Bidirectional crosstalk between Hypoxia-Inducible Factor and glucocorticoid signalling in zebrafish larvae. Plos Genetics. 2020;16(5).

9. Wang X, Zou Z, Yang Z, Jiang S, Lu Y, Wang D, et al. HIF 1 inhibits StAR transcription and testosterone synthesis in murine Leydig cells. J Mol Endocrinol. 2018.

10. Kowalewski MP, Gram A, Boos A. The role of hypoxia and HIF1alpha in the regulation of STAR-mediated steroidogenesis in granulosa cells. Mol Cell Endocrinol. 2015;401:35–44.

11. Yamashita K, Ito K, Endo J, Matsuhashi T, Katsumata Y, Yamamoto T, et al. Adrenal cortex hypoxia modulates aldosterone production in heart failure. Biochem Biophys Res Commun. 2020;524(1):184–9.

12. Madanecki P, Kapoor N, Bebok Z, Ochocka R, Collawn JF, Bartoszewski R. Regulation of angiogenesis by hypoxia: the role of microRNA. Cell Mol Biol Lett. 2013;18(1):47–57.

13. Komatsu S, Kitai H, Suzuki HI. Network Regulation of microRNA Biogenesis and Target Interaction. Cells. 2023;12(2).

14. Aparicio-Puerta E, Hirsch P, Schmartz GP, Fehlmann T, Keller V, Engel A, et al. isomiRdb: microRNA expression at isoform resolution. Nucleic Acids Research. 2023;51(D1):D179–D85.

15. Ong SG, Lee WH, Kodo K, Wu JC. MicroRNA-mediated regulation of differentiation and trans-differentiation in stem cells. Adv Drug Deliv Rev. 2015;88:3–15.

16. Zhao AH, Zeng Q, Xie XY, Zhou JN, Yue W, Li YL, et al. MicroRNA-125b Induces Cancer Cell Apoptosis Through Suppression of Bcl-2 Expression. J Genet Genomics. 2012;39(1):29–35.

17. Saliminejad K, Khorram Khorshid HR, Soleymani Fard S, Ghaffari SH. An overview of microRNAs: Biology, functions, therapeutics, and analysis methods. J Cell Physiol. 2019;234(5):5451–65.

18. Zaccagnini G, Greco S, Voellenkle C, Gaetano C, Martelli F. miR-210 hypoxamiR in Angiogenesis and Diabetes. Antioxid Redox Sign. 2022;36(10):685–706.

19. Seong M, Lee J, Kang H. Hypoxia-induced regulation of mTOR signaling by miR-7 targeting REDD1. Journal of cellular biochemistry. 2019;120(3):4523–32.

20. Bertero T, Rezzonico R, Pottier N, Mari B. Impact of MicroRNAs in the Cellular Response to Hypoxia. Int Rev Cel Mol Bio. 2017;333:91–158.

21. Hu Z, Shen WJ, Kraemer FB, Azhar S. Regulation of adrenal and ovarian steroidogenesis by miR-132. J Mol Endocrinol. 2017;59(3):269–83.

22. Azhar S, Dong D, Shen WJ, Hu Z, Kraemer FB. The role of miRNAs in regulating adrenal and gonadal steroidogenesis. J Mol Endocrinol. 2020;64(1):R21–R43.

23. Lai KP, Li JW, Tse ACK, Chan TF, Wu RSS. Hypoxia alters steroidogenesis in female marine medaka through miRNAs regulation. Aquatic Toxicology. 2016;172:1–8.

24. Nusrin S, Tong SK, Chaturvedi G, Wu RS, Giesy JP, Kong RY. Regulation of CYP11B1 and CYP11B2 steroidogenic genes by hypoxia-inducible miR-10b in H295R cells. Mar Pollut Bull. 2014;85(2):344–51.

25. Peitzsch M, Dekkers T, Haase M, Sweep FC, Quack I, Antoch G, et al. An LC-MS/MS method for steroid profiling during adrenal venous sampling for investigation of primary aldosteronism. J Steroid Biochem Mol Biol. 2015;145:75–84.

26. Bechmann N, Watts D, Steenblock C, Wallace PW, Schurmann A, Bornstein SR, et al. Adrenal Hormone Interactions and Metabolism: A Single Sample Multi-Omics Approach. Horm Metab Res. 2021;53(5):326–34.

27. Schindelin J, Arganda-Carreras I, Frise E, Kaynig V, Longair M, Pietzsch T, et al. Fiji: an open-source platform for biological-image analysis. Nature methods. 2012;9(7):676-82.

28. Pan Q, Kuang X, Cai S, Wang X, Du D, Wang J, et al. miR-132-3p priming enhances the effects of mesenchymal stromal cell-derived exosomes on ameliorating brain ischemic injury. Stem Cell Res Ther. 2020;11(1):260.

29. Olmo IG, Olmo RP, Gonçalves ANA, Pires RGW, Marques JT, Ribeiro FM. High-Throughput Sequencing of BACHD Mice Reveals Upregulation of Neuroprotective miRNAs at the Pre-Symptomatic Stage of Huntington’s Disease. Asn Neuro. 2021;13.

30. Han SR, Kang YH, Jeon H, Lee S, Park SJ, Song DY, et al. Differential Expression of miRNAs and Behavioral Change in the Cuprizone-Induced Demyelination Mouse Model. Int J Mol Sci. 2020;21(2).

31. Hyun J, Park J, Wang S, Kim J, Lee HH, Seo YS, et al. MicroRNA Expression Profiling in CCl(4)-Induced Liver Fibrosis of Mus musculus. Int J Mol Sci. 2016;17(6).

32. Zhang Y, Li CY, Guan C, Zhou B, Wang L, Yang CY, et al. MiR-181d-5p Targets KLF6 to Improve Ischemia/Reperfusion-Induced AKI Through Effects on Renal Function, Apoptosis, and Inflammation. Frontiers in Physiology. 2020;11.

33. Chen D, Lan G, Li R, Mei Y, Shui X, Gu X, et al. Melatonin ameliorates tau-related pathology via the miR-504-3p and CDK5 axis in Alzheimer’s disease. Transl Neurodegener. 2022;11(1):27.

34. Charania MA, Ayyadurai S, Ingersoll SA, Xiao B, Viennois E, Yan Y, et al. Intestinal epithelial CD98 synthesis specifically modulates expression of colonic microRNAs during colitis. Am J Physiol Gastrointest Liver Physiol. 2012;302(11):G1282–91.

35. Wang F, Qian J, Yang MY, He Y, Tao X, Shi YX, et al. MiR-6924-5p-rich exosomes derived from genetically modified Scleraxis-overexpressing PDGFRα(+) BMMSCs as novel nanotherapeutics for treating osteolysis during tendon-bone healing and improving healing strength. Biomaterials. 2021;279.

36. Mole DR, Blancher C, Copley RR, Pollard PJ, Gleadle JM, Ragoussis J, et al. Genome-wide association of hypoxia-inducible factor (HIF)-1alpha and HIF-2alpha DNA binding with expression profiling of hypoxia-inducible transcripts. J Biol Chem. 2009;284(25):16767–75.

37. Schodel J, Oikonomopoulos S, Ragoussis J, Pugh CW, Ratcliffe PJ, Mole DR. High-resolution genome-wide mapping of HIF-binding sites by ChIP-seq. Blood. 2011;117(23):e207–17.

38. Stocco DM. StAR protein and the regulation of steroid hormone biosynthesis. Annu Rev Physiol. 2001;63:193–213.

39. Hu Z, Shen WJ, Cortez Y, Tang X, Liu LF, Kraemer FB, et al. Hormonal regulation of microRNA expression in steroid producing cells of the ovary, testis and adrenal gland. PLoS One. 2013;8(10):e78040.

40. Takeda Y, Demura M, Kometani M, Karashima S, Yoneda T, Takeda Y. Molecular and Epigenetic Control of Aldosterone Synthase, CYP11B2 and 11-Hydroxylase, CYP11B1. Int J Mol Sci. 2023;24(6).

41. Filipowicz W, Bhattacharyya SN, Sonenberg N. Mechanisms of post-transcriptional regulation by microRNAs: are the answers in sight? Nat Rev Genet. 2008;9(2):102–14.

42. Afonso-Grunz F, Muller S. Principles of miRNA-mRNA interactions: beyond sequence complementarity. Cell Mol Life Sci. 2015;72(16):3127–41.

43. Kulshreshtha R, Davuluri RV, Calin GA, Ivan M. A microRNA component of the hypoxic response. Cell Death Differ. 2008;15(4):667–71.

44. Donker RB, Mouillet JF, Nelson DM, Sadovsky Y. The expression of Argonaute2 and related microRNA biogenesis proteins in normal and hypoxic trophoblasts. Mol Hum Reprod. 2007;13(4):273–9.

45. Nusrin S, Tong SKH, Chaturvedi G, Wu RSS, Giesy JP, Kong RYC. Regulation of CYP11B1 and CYP11B2 steroidogenic genes by hypoxia-inducible miR-10b in H295R cells. Marine Pollution Bulletin. 2014;85(2):344–51.

46. Chaurasiya V, Kumari S, Onteru SK, Singh D. miR-326 down-regulate expression and estradiol-17b production in buffalo granulosa cells through CREB and C/EBP-β. J Steroid Biochem. 2020;199.

47. Liu D, Zou S, Li G, Zhang Q, Chen C, Li C, et al. Downregulation of Uncoupling Protein 2(UCP2) Mediated by MicroRNA-762 Confers Cardioprotection and Participates in the Regulation of Dynamic Mitochondrial Homeostasis of Dynamin Related Protein1 (DRP1) After Myocardial Infarction in Mice. Front Cardiovasc Med. 2021;8:764064.

48. Zhu WJ, Huang HH, Feng YF, Zhan L, Yang J, Zhu L, et al. Hypoxia-induced miR-9 expression promotes ovarian cancer progression via activating PI3K/AKT/mTOR/GSK3beta signaling pathway. Neoplasma. 2023;70(2):216–28.

49. Liao Y, Li H, Cao H, Dong Y, Gao L, Liu Z, et al. Therapeutic silencing miR-146b-5p improves cardiac remodeling in a porcine model of myocardial infarction by modulating the wound reparative phenotype. Protein Cell. 2021;12(3):194–212.

50. Yu RM, Chaturvedi G, Tong SK, Nusrin S, Giesy JP, Wu RS, et al. Evidence for microRNA-mediated regulation of steroidogenesis by hypoxia. Environ Sci Technol. 2015;49(2):1138–47.

51. Vadysirisack DD, Ellisen LW. mTOR activity under hypoxia. Methods in molecular biology. 2012;821:45–58.

52. Uniacke J, Holterman CE, Lachance G, Franovic A, Jacob MD, Fabian MR, et al. An oxygen-regulated switch in the protein synthesis machinery. Nature. 2012;486(7401):126-9.

