## Supplementary figures and images for "HIF1α controls steroidogenesis under acute hypoxic stress"

### Supplementary Figures

#### Supplementary Figure 1

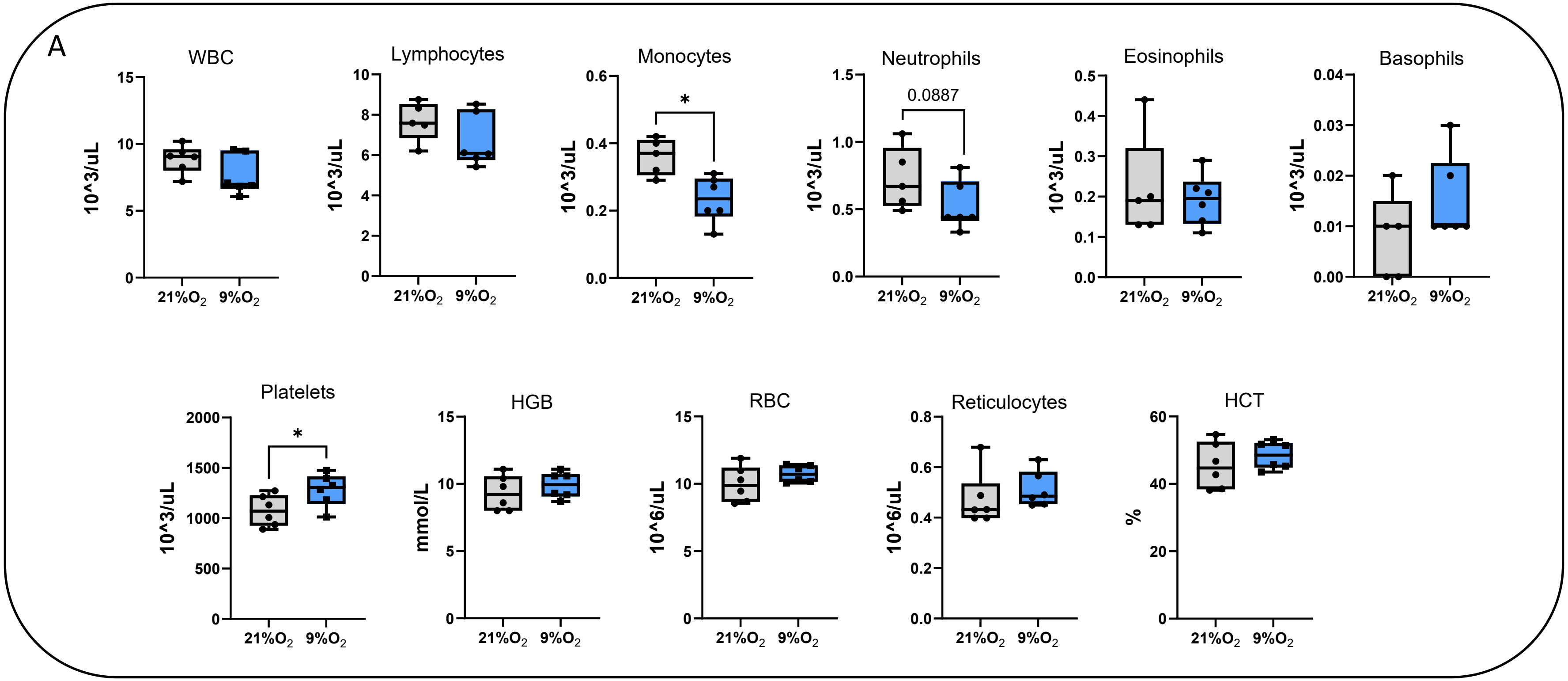


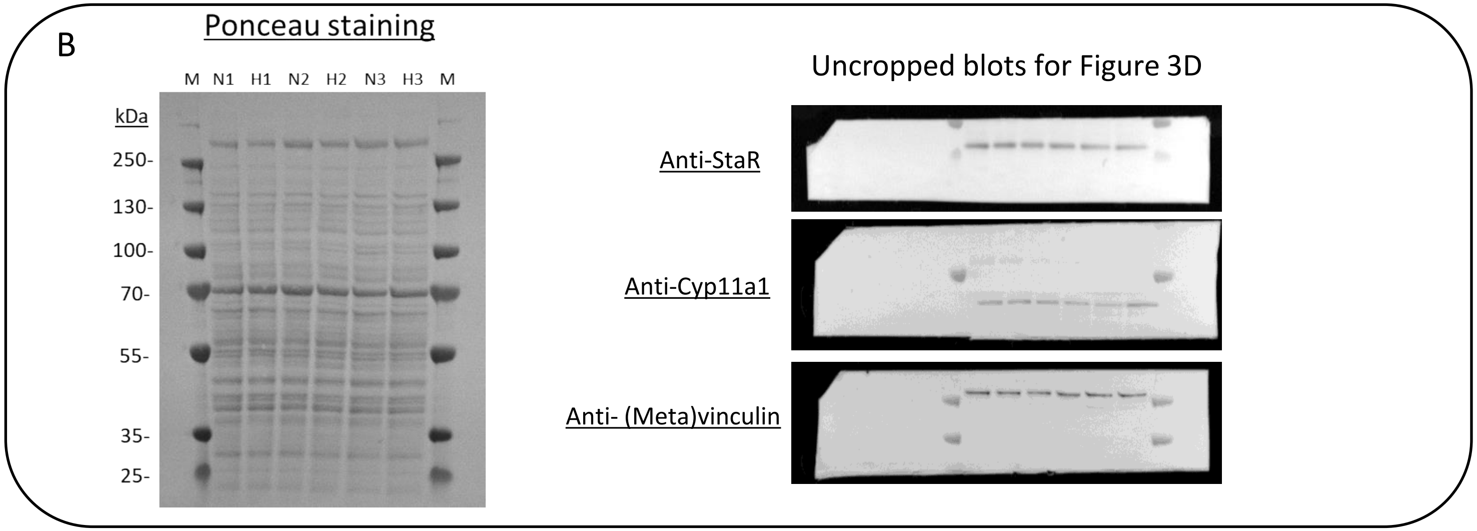

#### Supplementary Figure 2

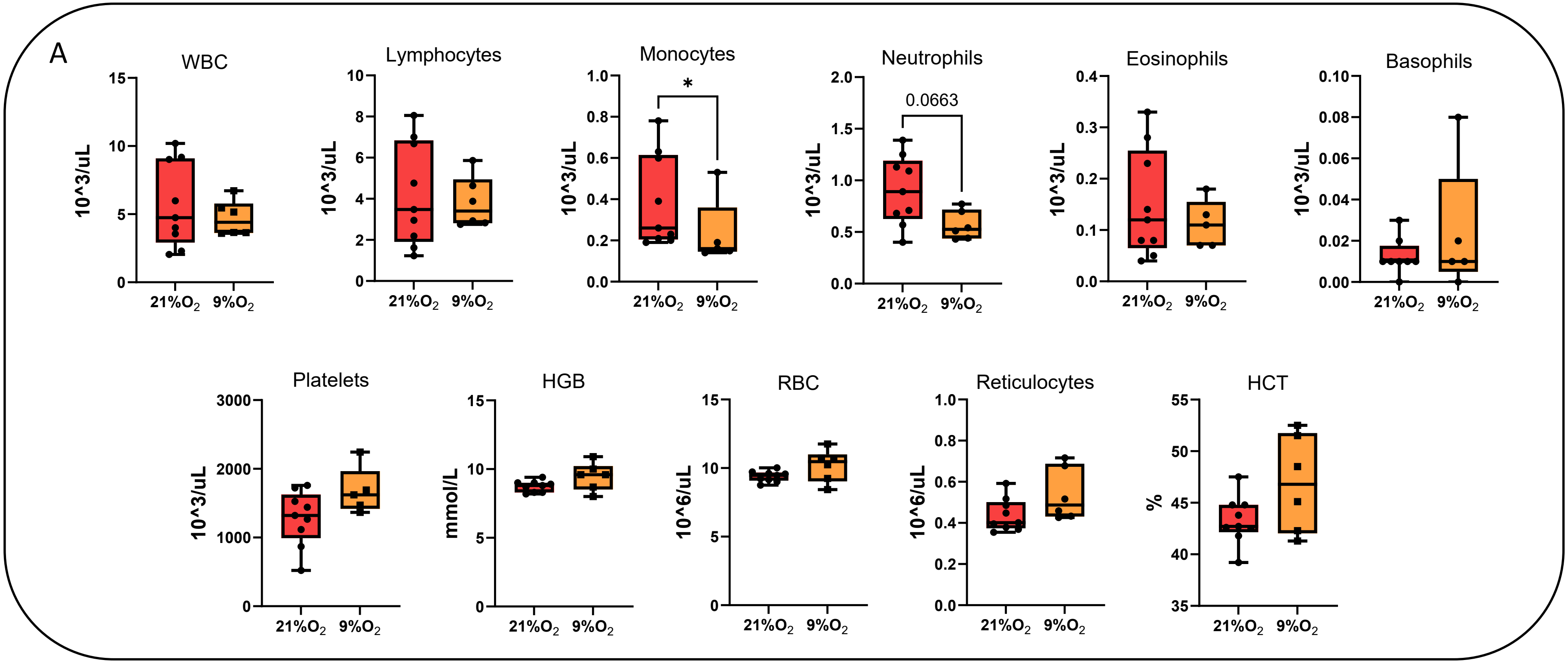


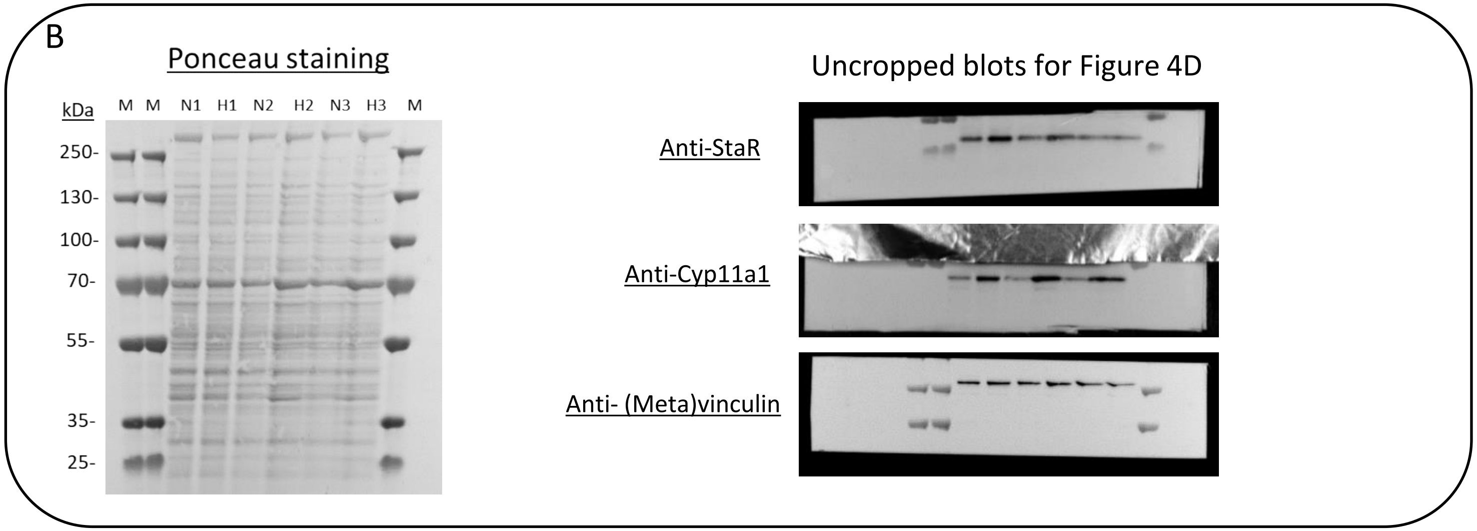
